# With great power comes great responsibility: an analysis of sustainable forest management quantitative indicators in the DPSIR framework

**DOI:** 10.1101/2021.02.11.430737

**Authors:** Y. Paillet, T. Campagnaro, S. Burrascano, M. Gosselin, J. Ballweg, F. Chianucci, J. Dorioz, J. Marsaud, L. Maciejewski, T. Sitzia, G. Vacchiano

## Abstract

The monitoring of environmental policies in Europe has taken place since the 1980s and still remains a challenge for decision- and policy-making. For forests, it is concretized through the publication of a State Of Europe’s Forests every five years, the last report just been released. However, the process lacks a clear analytical framework and appears limited to orient and truly assess sustainable management of European forests. We classified the 34 quantitative sustainable forest management indicators in the Driver-Pressure-State-Impact-Response (DPSIR) framework to analyse gaps in the process. In addition, we classified biodiversity-related indicators in the simpler Pressure-State-Response (PSR) framework. We showed that most of the sustainable forest management indicators assess the state of European forests, but almost half could be classified in another DPSIR category. For biodiversity, most indicators describe pressures, while direct taxonomic state indicators are very few. Our expert-based classification show that sustainable forest management indicators are unbalanced regarding the DPSIR framework. However, completing this framework with other indicators would help to have a better view and more relevant tools for decision-making. The results for biodiversity were comparable, but we showed that some indicators from other criteria than the one dedicated to biodiversity could also help understanding threats and actions concerning it. Such classification helps in the decision process, but is not sufficient to fully support policy initiative. In particular, the next step would be to better understand the links between DPSIR and PSR categories.

## Introduction

Monitoring the performance of sectorial policies has long been a challenge for policy makers, searching for legitimacy and interest of a large panel of stakeholders (Marsh, 1999). They might be interested in assessing whether quantitative goals have been achieved, or if an investment has produced the expected effects, or more generally how the various taxes collected by a given entity have been re-invested in public or private actions (Butchart et al., 2010). Hence the challenge resides in producing fully transparent, reproducible and accessible monitoring of environmental policies targeting biodiversity conservation or sustainable forest management (Milieu Ltd et al., 2016, Puumalainen et al., 2003). The definition of sustainable forest management recently shifted from the consolidated one: “stewardship and use of forests and forest lands in such a way, and at a rate, that maintains their biodiversity, productivity, regeneration capacity, vitality and their potential to fulfil, now and in the future, relevant ecological, economic and social functions, at local, national, and global levels, and that does not cause damage to other ecosystems” (FOREST EUROPE, 2020) to “practices and uses of forests and forest land that contribute to enhancing biodiversity or to halting or preventing the degradation of ecosystems, deforestation and habitat loss”. This latter definition takes into account the former one in the regulation known as “Sustainable Finance Taxonomy” (EU Regulation 2020/852). Such shifts means a substantial change from the association of sustainability to multifunctionality towards a strict focus on biodiversity and habitat conservation, and requires to rethink the methods of monitoring the levels of achievement of sustainable forest management.

Monitoring of environmental policies in Europe has taken place since the 1980s, and is concretized through the publication of the State Of Europe’s Forests every five years (FOREST EUROPE, 2020). This process has consisted in building a Pan-European criteria and indicators framework dedicated to the subject at stake. In this report, 34 quantitative indicators are distributed between six criteria covering the whole panel of objectives related to sustainable forest management (Table 1, FOREST EUROPE, 2020). These indicators have been mostly chosen using pre-existing data (notably National Forest Inventories, e.g. Chirici et al., 2012 for biodiversity, and the recent Heym et al., 2021) and are more consensus-oriented rather than question-driven, as identified in many other processes aiming at monitoring environmental policies (Scholes et al., 2012). However, policy and management assessments remain relatively limited and addressed by this process (see e.g. Feest, 2013 in the case of the Streamlining European Biodiversity Indicators - SEBI), notably due to the lack of explicit analytical framework. This need for better, more easily monitorable indicators is also apparent from efforts currently underway to use remote sensing as a biodiversity monitoring method (e.g., the GEOBON framework and “essential biodiversity variables”, Scholes et al., 2012, Pereira et al., 2013). Despite interesting use of multi-criteria analyses to improve effectiveness of sustainable forest management indicators (Vacik et al., 2007, Baycheva-Merger and Wolfslehner, 2016) no explicit overall analysis has been attempted to date. Rather, indicators have been generally considered separately and independently (Gao et al., 2015), assuming that they can effectively and comprehensively describe the trend of the criterion to which they belong (see however Wolfslehner and Vacik, 2011)). For example, in the State of Europe’s Forests, the criterion 4 dealing with biodiversity is composed of ten indicators that are neither weighted, ranked, nor analyzed in an integrated or transversal manner with respect to the problematic they are supposed to answer. Namely, within the criterion “Maintenance, conservation and appropriate enhancement of biological diversity in forest ecosystems”, biodiversity is assessed through what one could call indirect (i.e. non-taxonomic) indicators or proxies like deadwood volume and very few direct taxonomic data, nor, for that matter, indices of taxonomic or functional diversity are used (Lier et al., 2013, Paillet et al., 2013). In fact, the monitoring process of the EU ecosystems condition notes that current forest indicators are either limited in time, spatial scale or are relative to few dimensions (Maes et al., 2018). The reasons behind these limitations are the high costs and the wide range of skills needed to directly measure overall (i.e., multi-taxonomic) forest biodiversity. The multidimensional space of biological diversity encompassing enormously different organisms at highly different spatial scales makes a holistic assessment of the overall forest biodiversity unlikely. This, in turn, hinders the implementation of sustainable forest management strategies to balance biodiversity conservation targets, maintenance of the health and productive capacity of forest ecosystems and their role in watershed and carbon cycle.

**Table 1:**
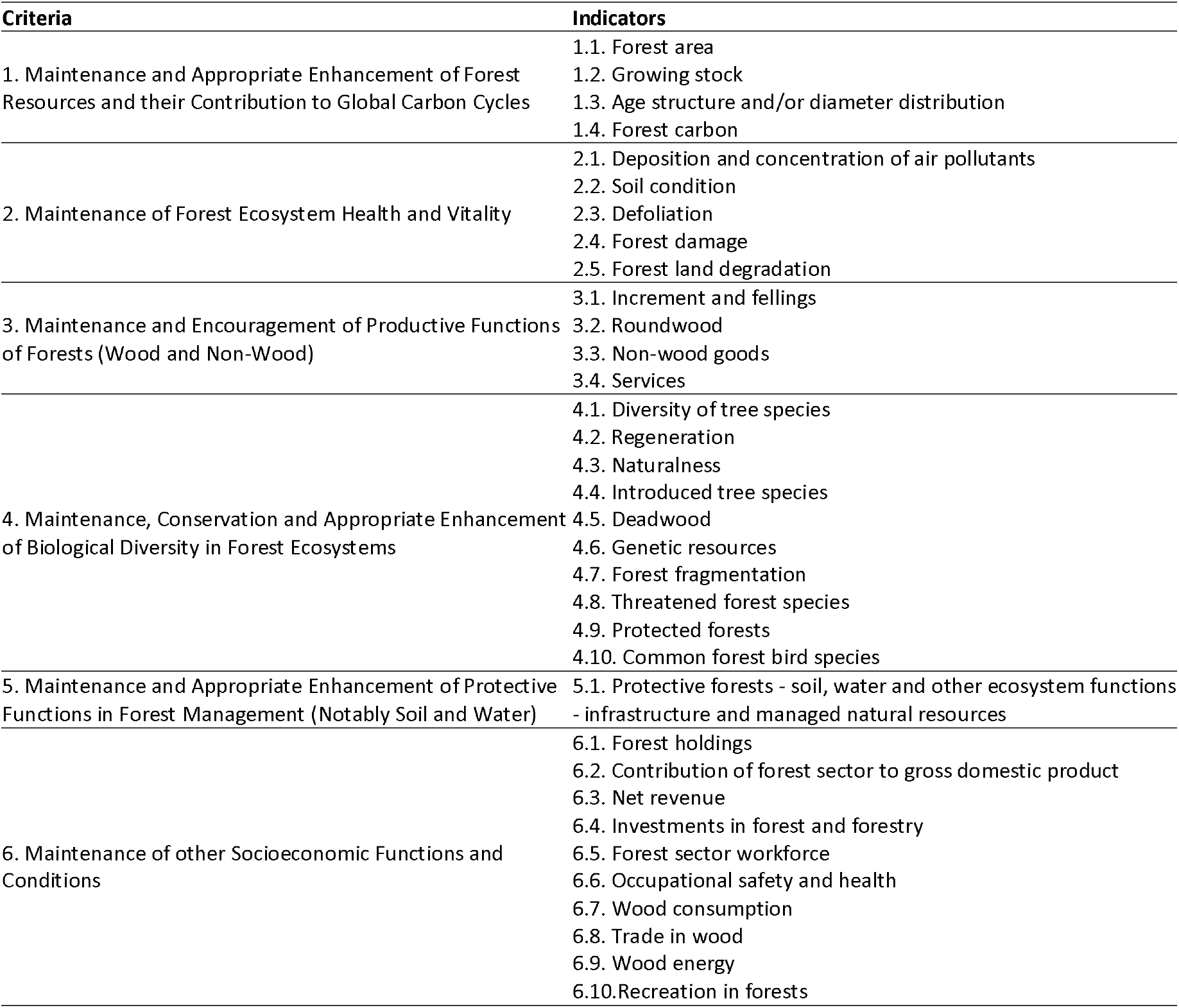
Quantitative criteria and indicators of Sustainable Forest management (FOREST EUROPE 2020).

Various analytical frameworks have been developed to assess the link between society and environment as well as associated issues. The Driver-Pressure-State-Impact-Response (DPSIR) is the most famous and was notably adopted by the European Environment Agency (EEA 1999). According to the DPSIR framework, there is a chain of assumed causal links: “driving forces” (economic sectors, human activities) exert “pressures” (emissions, waste, etc) on “states” of ecosystems (physical, chemical and biological) that bear “impacts” on human well-being, and eventually lead to political “responses” (e.g. prioritization, target setting, EEA, 1999). The DPSIR framework was first conceptualized as a flexible framework aimed to assist decision-making (Bell, 2012, Tscherning et al., 2012), then was used to appropriately structure and organize indicators systems (Tscherning et al., 2012). DPSIR was originally developed by the Organisation for Economic Co-operation and Development (OECD, 1994) and used by the United Nations (UNEP, 1997, UNEP, 2007) and European Environmental Agency (EEA, 1999) to link human activities to the environment. In the case of the European Environment Agency, the Streamlining European Biodiversity Indicators are based on – or developed within – the DPSIR, but no analysis linking the different categories of the framework has been explicitly presented yet (Feest, 2013). More surprisingly, in the case of sustainable forest management criteria and indicators, few *ex-post* assessment of FOREST EUROPE’s indicators has been attempted to date, except locally and based on local stakeholders involvement (e.g. Vacik et al., 2007, Scriban et al., 2019).

In this context, we aimed to gauge whether sustainable forest management and biodiversity indicators of FOREST EUROPE (2020) were usable to answer policy questions. For this reason, we first classified the existing sustainable forest management indicators (FOREST EUROPE, 2020) within the DPSIR framework using referenced and reproducible methods in order to identify the strengths and weaknesses of the current indicator system as well as its completeness with respect to the DPSIR framework. We secondly focused on the biodiversity criterion and classified its indicators within a restricted Pressure-State-Response (PSR) framework.

## Methods

### Use of the DPSIR framework

We adopted the European Environmental Agency and the Organisation for Economic Co-operation and Development definitions of the DPSIR and PSR frameworks, respectively, to analyze environmental issues as mentioned above (Figure 1). In the DPSIR conceptual framework, categories are defined as follows (adapted from EEA, 1999):

**Figure 1:**
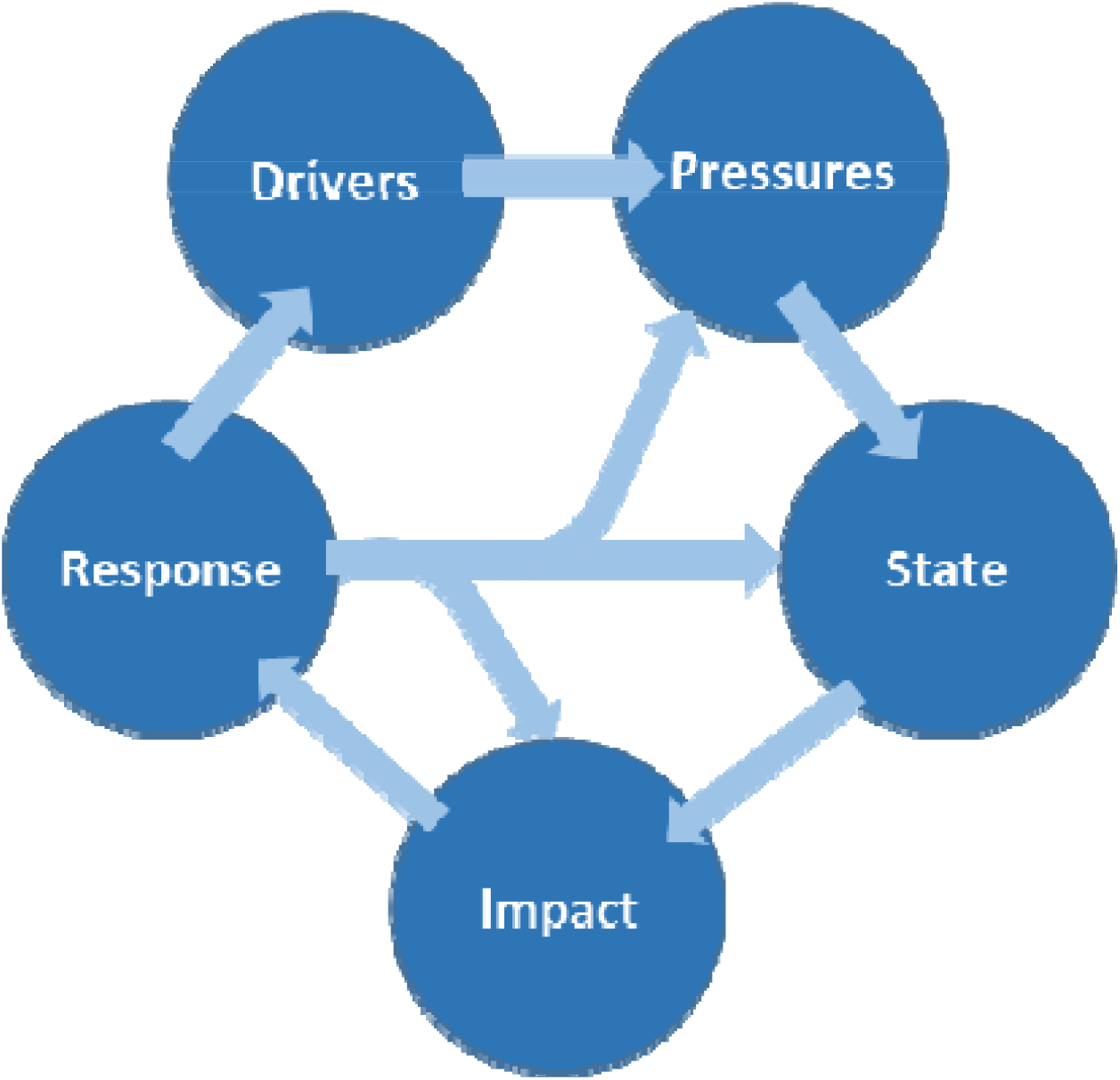
Schematic representation of the DPSIR framework

– <u>Drivers</u> are the demographic, social and economic developments within a given society and their corresponding changes in life styles, overall levels of consumption and production patterns;
– <u>Pressures</u> are the direct causes of degradation of the object assessed and, as such, have generally a negative connotation. We classified them according to the HIPOC model refers to the five main threats to biodiversity which are Habitat conversion and degradation, Invasion of alien species, Pollution and eutrophication, Over-exploitation and unsustainable use, Climate change mentioned in Maes et al. (2018). Usually, changes linked with these pressures are unwanted and seen as negative (e.g. damage, degradation) although a decrease in the intensity of a pressure may result in an improvement of the condition of an ecosystem (Maes et al., 2018). The pressures exerted by society, such as habitat destruction, may directly or indirectly affect ecosystem conditions;
– <u>State</u> stands for the condition of the abiotic and biotic components of the ecosystems in a given area. It consists of quantities and qualities of physical, chemical or biological variables. Biological variables concern the condition at the ecosystem, habitat, species, community, or genetic levels (according to the definition of the Convention on Biological Diversity);
– Changes in the quality and functioning of the ecosystem have an <u>impact</u> on the welfare or well-being of humans through the provision of ecosystem services. Biodiversity and more generally the environment contributes to ecosystem goods and services, which in turn directly or indirectly cause human social or economic benefits, or have the potential to do so in the future. The definition used here derives from the seminal version of the MEA (Millenium Ecosystem Assessment, 2005) in the Economics of Ecosystems and Biodiversity report (TEEB, 2010). It identifies the following types of services: provisioning, regulating and cultural services (supporting services, initially included in the MEA, are more seen as functions). Environmental impacts, such as considered in the EU Framework Water Directive (e.g. Borja et al., 2006), were not classified as such here;
– <u>Responses</u> are actions taken by governmental and non-governmental institutions or parties (groups, individuals, governments) to prevent, mitigate, compensate for, improve or adapt to, changes in the state of the environment by seeking to (i) control drivers or pressures through regulation, prevention, or mitigation; (ii) directly maintain or restore the state of the environment; or (iii) deliberately “do nothing”. The definition and update of the EU Forest Strategy (European Commission, 2013), as well as the forestry measures included in the Common Agricultural Policy or the established conservation measures of the Habitat Directive may represent the responses to the impact reported on forest ecosystems.

The interest of the DPSIR framework resides in its circular interlinked pattern (Figure 1) that stands for its causal chain nature. In other words, for a given subject, it is theoretically possible to link each component of the scheme with another along the causal chain. This requires the specification of a precise object and formulation of explicit question(s) to be answered by a dedicated indicator set (Bubb et al., 2010, Niemeijer and de Groot, 2008, Gao et al., 2015).

### Objects and questions

We defined the sustainable forest management and forest biodiversity policy questions to be answered and the indicanda, i.e. the objects to be assessed by each indicator (Heink and Kowarik, 2010a), as follows:

1. Sustainable Forest Management indicators: are European forests managed in a sustainable way, according to the definition provided by FOREST EUROPE (2020)? The forest socio-ecosystem is the indicandum.
2. Biodiversity: is (forest) biodiversity maintained, conserved, improved or restored in European forests? Biodiversity, as defined by the Secretariat of the Convention on Biological Diversity (2006) is the indicandum.

To analyse the relevance of sustainable forest management indicators to answer the question (1), we adopted the entire DPSIR framework. For question (2), we restricted the analysis to the simpler PSR framework as the links between drivers and pressures, as well as links between state and impacts, may be subject to unsolvable debates due to lack of quantitative knowledge concerning the role of biodiversity in ecosystem functioning and services (Duncan et al., 2015, Isbell et al., 2018). In addition, drivers and impacts were likely to be the same as those potentially affecting forests in the first analysis.

### Approach adopted

Regarding the definitions above, we deliberately adopted a precise and stable definition of the DPSIR (PSR) framework, contrary to the classical view of DPSIR as a flexible framework (see Gari et al., 2015 and references therein). Our approach is essentially based on the expertise of the authors but also considers scientific literature on the subject (Wolfslehner, 2007). By clearly expliciting the classifications we used and delineating the boundaries and potential limitations of our methods (Heink and Kowarik, 2010a), we expected it to be reproducible and relevant (Failing and Gregory, 2003).

For a given indicandum, we classified each indicator in one, and only one, DPSIR or PSR category. For pressure indicators, based on expertise of the authors, we specified the direction and magnitude of the effect caused by a given increase in the indicator value on a categorical scale (++, +, equivocal: +/-, - and --) and the part of biodiversity affected by a given pressure (e.g. forest specialist or generalists species). We also specified the pressure type in the HIPOC classification. For sustainable forest management indicators, we considered only those describing physical, chemical or biological conditions of forests (direct indicators) as state indicators. Similarly, for biodiversity indicators, we considered only taxonomic indicators (direct indicators) as state indicators, while proxies (indirect indicators) were classified as pressure indicators (see discussion). In the case of biodiversity, we not only analyzed the indicators of the biodiversity criterion (number 4 in the FOREST EUROPE framework), but also considered the indicators from all the other five criteria that may have an influence on biodiversity (Table 1). For impact indicators, we specified the broad category of ecosystem service being impacted as defined by the TEEB (2010).

## Results

### Classification of Forest Europe’s sustainable forest management indicators (indicandum = forests)

First, all Forest Europe’s indicators were classified into one DPSIR category without ambiguity (Table 2, Figure 2). The large majority of indicators were easily classified in one of the sub-categories we used, except for two exceptions:

**Table 2:** Classification of Forest Europe’s sustainable forest management indicators in the DPSIR framework. Each indicator was classified in only one category. + and – signs relate to the intensity of the effect on the forest ecosystem linked with an increase in the individual indicator value, based on the author’s expertise: -- strong negative, - negative, +/- equivocal, + positive, ++ strong positive effects.

| Indicators | Driver | Pressure<br>(HIPOC category) | State | Impact<br>(Ecosystem Service<br>category) | Response |
| --- | --- | --- | --- | --- | --- |
| 1.1. Forest area |  |  | X |  |  |
| 1.2. Growing stock |  |  | X |  |  |
| 1.3. Age structure and/or diameter distribution |  |  | X |  |  |
| 1.4. Forest carbon |  |  | X | Regulating |  |
| 2.1. Deposition and concentration of air pollutants |  | Pollution (--) |  |  |  |
| 2.2. Soil condition |  |  | X |  |  |
| 2.3. Defoliation |  |  | X |  |  |
| 2.4. Forest damage |  |  | X |  |  |
| 2.5. Forest land degradation |  | Habitat loss (--) |  |  |  |
| 3.1. Increment and felling |  | Over exploitation<br>(felling, +/-) | X<br>(increment) |  |  |
| 3.2. Roundwood |  |  |  | Provisioning |  |
| 3.3. Non-wood goods |  |  |  | Provisioning |  |
| 3.4. Services (value) |  |  |  | Provisioning |  |
| 4.1. Diversity of tree species |  |  | X |  |  |
| 4.2. Regeneration |  |  | X |  |  |
| 4.3. Naturalness |  |  | X |  |  |
| 4.4. Introduced tree species |  |  | X |  |  |
| 4.5. Deadwood |  |  | X |  |  |
| 4.6. Genetic resources |  |  |  |  | X |
| 4.7. Forest fragmentation |  |  | X |  |  |
| 4.8. Threatened forest species |  |  | X |  |  |
| 4.9. Protected forests |  |  |  |  | X |
| 4.10. Common forest bird species |  |  | X |  |  |
| 5.1. Protective forests |  |  |  | Regulating |  |
| 6.1. Forest holdings | X |  |  |  |  |
| 6.2. Contribution of forest sector to gross domestic product |  |  |  | Provisioning |  |
| 6.3. Net revenue |  |  |  | Provisioning |  |
| 6.4. Investments in forests and forestry |  |  |  |  | X |
| 6.5. Forest sector workforces |  |  |  | Provisioning |  |
| 6.6. Occupational safety and health |  |  |  | Regulating |  |
| 6.7. Wood consumption | X |  |  |  |  |
| 6.8. Trade in wood | X |  |  |  |  |
| 6.9. Wood energy |  |  |  | Provisioning<br>Regulating |  |
| 6.10. Recreation in forests |  |  |  | Cultural |  |
| **Total** | **3** | **3** | **16** | **11** | **3** |

**Figure 2:**
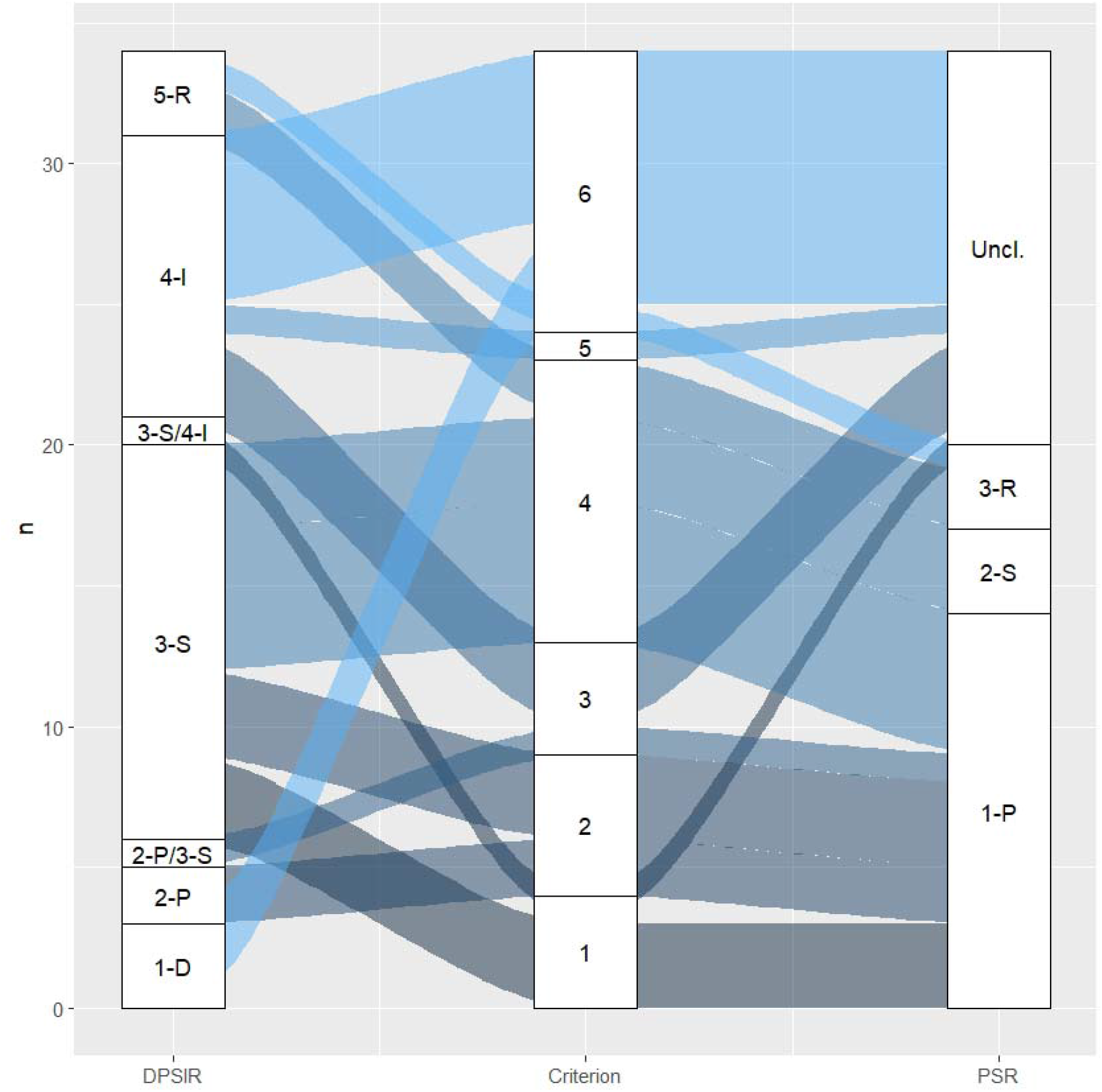
Classification of the sustainable forests management quantitative indicators in the Driver-Pressure-State-Impact-Response (DPSIR, numbered from 1 to 5) framework (left) and the indicators linked with biodiversity in the PSR (Pressure-State-Response, numbered from 1 to 3) framework (right). Numbers refer to the criteria of sustainable forest management (FOREST EUROPE, 2020, see Table 1). Uncl. corresponds to “Unclassified” indicators in the PSR framework.

– The carbon stock indicator (1.4) that could be either a state of forests or an impact (regulating) indicator (see below);
– Increment and fellings (3.1) describes two different processes: “increment” was classified as state, while “felling” was classified as pressure.

Note also that the wood energy indicator (6.9) indicate either a provisioning or a regulating service, since it also allows mitigation via substitution of fossil fuels.

Globally, sustainable forest management indicators were inequitably distributed in the DPSIR classification (Figure 2a): only three driver and three pressure indicators were identified. The majority of indicators related to state of the forest ecosystem (16) or impacts in terms of ecosystem services (9). More marginally, three indicators related to a society’s response in terms of sustainable forest management.

The three driver indicators concerned forest holdings, wood consumption and trade in wood.

The three pressure indicators concerned pollution, habitat loss and over-harvesting with negative effects of air pollutants (2.1) and land degradation (2.5), while the effect of fellings (3.1) was equivocal (ie. either negative or positive depending on the part of the ecosystem assessed and management type).

Sixteen indicators were considered as state indicators. Most of them described biological forest characteristics, i.e. structure and composition, but not functioning except for soil conditions (abiotic physical and chemical attributes). We classified the indicator related to carbon stocks (1.4) as a state indicator since it describes the quantity of carbon in forests but could also be interpreted in terms the capacity or flux, that would better correspond more to an impact indicator (regulating service).

Eleven indicators described impacts in terms of ecosystem services. In addition to forest carbon (1.4., regulating service), six were indicating provisioning services, two regulating services, and one cultural services. As mentioned above only one was classified in two categories of services (6.9 Wood energy).

Finally, only three indicators were considered as response indicators, either of institutional nature (protected forests) or reflecting more incentives and values potentially beneficial to biodiversity (conservation of genetic resources, investments in forests and forestry).

### Classification of Forest Europe’s biodiversity-related indicators (indicandum = biodiversity)

Nearly two thirds (twenty-one) of the thirty-four indicators were considered as related to biodiversity in a more or less explicit way Table3, (Figure 2). Evidently, all the indicators of the biodiversity criterion were included in this group, while most of the others were issued from criteria 1 and 2. Two thirds (fourteen) out of the twenty-two indicators were classified as pressure indicators, mostly related to habitat loss (or gains) and over-harvesting, the difference between which was not always clear (e.g. for age structure, regeneration, see Table 3). Based on the author’s expertise, five indicators translated into positive effects on either forest generalists and/or specialists (three strong, the others moderate). Six others translated into negative effects while two had equivocal effects (three strong, the others moderate). Some pressure indicators concerned only certain ecological groups such as saproxylics (indicator 4.5 Deadwood) or forest core and dispersal limited species (4.7 Landscape patterns). As for the classification of sustainable forest management indicators in the DPSIR, some indicators comprising several dimensions had contrasting effects on biodiversity, e.g. naturalness (4.3.) could have either positive (natural forests) or negative (plantations) effects on saproxylic species. The indicator relating to fellings, that could be seen as a pressure on biodiversity, was not classified since it was too tricky to affect either a HIPOC category or an effect on biodiversity, that depends on intensity of fellings and the part of the biodiversity considered.

**Table 3:**
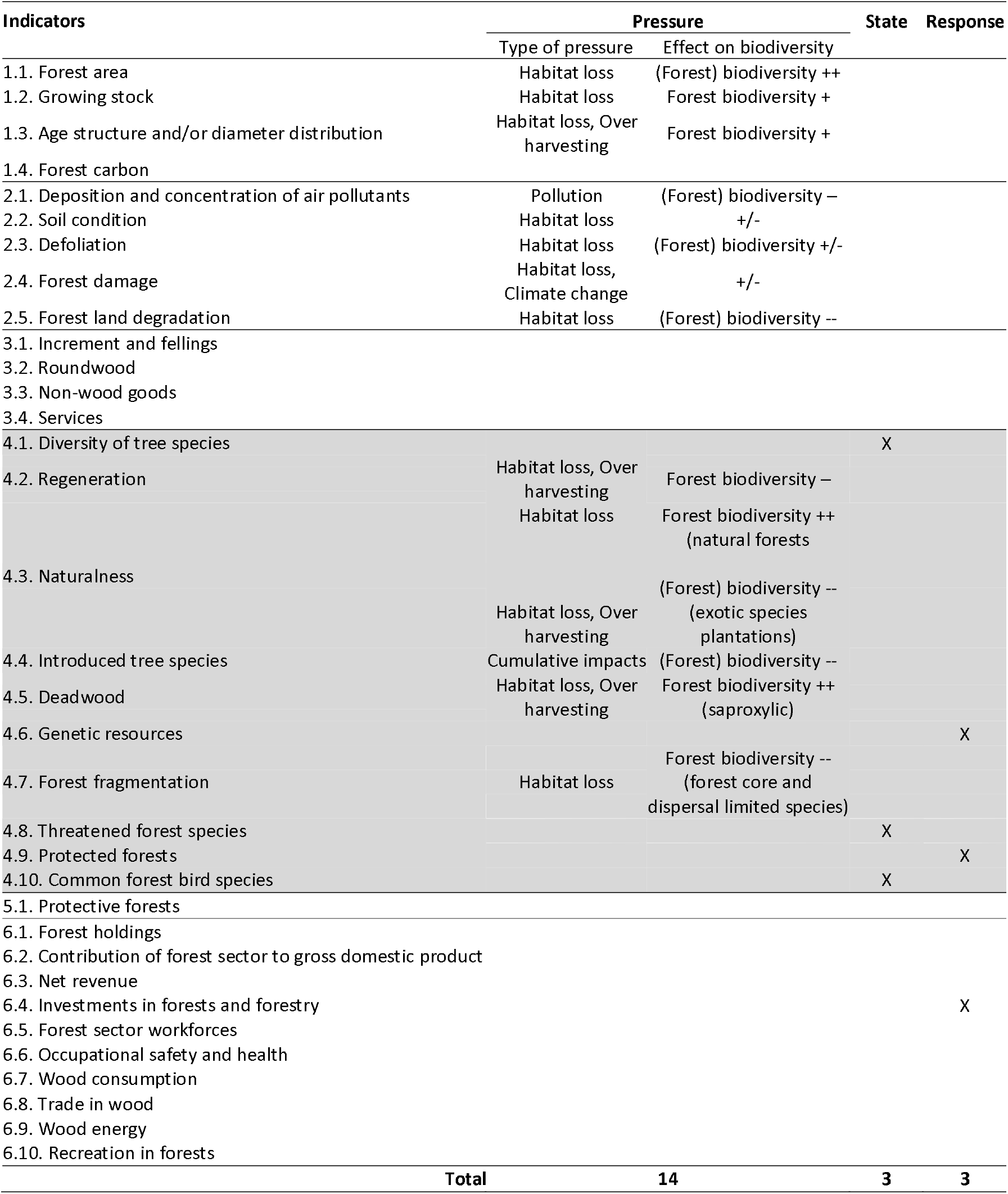
Classification of sustainable forest management indicators in the PSR framework with respect to biodiversity. All indicators that related to biodiversity outside the dedicated criterion (4) were also classified. Components of biodiversity concerned by pressures: (Forest) biodiversity = both generalist and specialist forest species; Forest biodiversity= specialist forest species. + and – signs relate to the intensity of the effect on biodiversity linked with an increase in the individual indicator value, based on the author’s expertise: -- strong negative, - negative, +/- equivocal, + positive, ++ strong positive effects.

The most striking result concerned state (taxonomic) indicators that were only three concerning species diversity (4.1), threatened forest species (4.8) and common forest birds (4.10).

Finally, four indicators were classified as response, most of them relating to forest management tools that could be beneficial to biodiversity (forests protected for biodiversity or protection forests), but also to other values (Investments in forests and forestry).

All the other indicators were not classified because they had very weak (or no) direct links with biodiversity, especially most of those issued from the criterion 6.

## Discussion

By classifying Forest Europe’s indicators in the DPSIR framework – resp. biodiversity-related indicators in the PSR framework – we provide an insight of their power and limitations to assess sustainable forest management. Explicit definitions the indicanda “forest” and “biodiversity”, ie the objects to be assessed, and mainstream classifications allowed us to analyse gaps and recommendations to fill them in. However, the system adopted remains imperfect with blurred lines between some categories, and limitations in the DPSIR and PSR frameworks.

### Analysing sustainable forest management and biodiversity indicators through the DPSIR

Numerous environmental indicators suffer from a lack of precision in the definition of their indicanda (Gao et al., 2015, Niemeijer and de Groot, 2008, Paillet et al., 2013). Sustainable Forest Management indicators used at the European level are no exception to the rule, although they are supposed to describe the state of Europe’s forest (FOREST EUROPE, 2020), no explicit links with indicanda within an analytic framework has been attempted so far, which limits their use for policy assessment and orientation (Feest, 2013). As obvious as it seems, clearly defining the indicandum when validating or using indicators is a step that is too often neglected (Gao et al., 2015, Gari et al., 2015, Heink and Kowarik, 2010a). By choosing forest ecosystem and biodiversity as indicanda in two different analyses using the DPSIR and PSR frameworks, respectively, we first clarified the purpose of sustainable forest management and biodiversity indicators. We showed that changing the indicandum also modified the way we interpret indicators. Indeed, the very same indicator, e.g. deadwood, may assess the state of the forest ecosystem, but becomes a pressure indicator when it comes to assessing biodiversity of forest dwelling species (Tables 1 & 2). Of course this depends on the choice we made to consider only direct taxonomic indicators as state indicators of biodiversity, while other, habitat-related indicators, were classified as pressures, but could also be considered as ecosystem-level biodiversity state indicators describing the quality of the habitat (Scholes et al., 2012). Nevertheless, we assume that this choice clarifies the use of biodiversity indicators for policy and management (see below). In terms of assessment, since environmental indicators should reveal more than their measurements *per se* (Blauvelt, 2014), it means that a clear definition of the object for which the indicator is identified is crucial to drive decision processes and policymaking. This includes prioritising objectives, identifying and choosing alternatives that satisfy the objectives, and implementing the policy decisions. Indeed, a given indicator may eventually show misleading results depending on the chosen indicandum. It also conditions the way scientists, policy makers, stakeholders and the general public communicate (Blauvelt, 2014). That said, our classifications show that sustainable forest management and biodiversity indicators are relatively incomplete and do not provide an overall vision of the situation regarding the DPSIR (PSR) framework.

Concerning sustainable forest management indicators distributed in the DPSIR classification, we showed that driver, pressure and response indicators are almost absent from the process: 3 each, out of 34, which summed up represent merely a fourth of the total number of indicators. Indeed, the indicators of sustainable forest management mainly assess the state of forest as exemplified by the name of the dedicated publication entitled “State of Europe’s Forest” (FOREST EUROPE, 2020), and its impact on human well-being through what we classified as services. On the other hand, we still identified drivers and responses in the process so that sustainable forest management indicators do not only refer to the “State of Forests”. Such results may be explained by the fact that the data used to calculate these indicators are mostly issued by National Forest Inventories that basically describe composition and structure of the ecosystem (Tomppo et al., 2010, Alberdi et al., 2019, Vidal et al., 2016). Other indicators are issued by market-based studies and databases, which allow an interpretation in terms of insertion of forest products and values in the global economy (Linser et al., 2018). However, when replaced within an analytical framework such as the DPSIR, the lack of driver, but mostly of pressure, indicators does not allow to have a complete view of the processes that drive the forest ecosystem. For example, no indicator of climate change is included in the indicator set yet, despite the growing importance of this pressure on the forest ecosystem (Lindner et al., 2010). Therefore, it is likely that sustainable forest management indicators are of relatively poor value to support decision-making processes that aim at reducing pressure on forests through the use of appropriate institutional commitments (namely responses, that are also poorly assessed by the process) and are difficult to analyse in a transversal way.

Concerning biodiversity-related indicators, the situation is different. When classified in the PSR framework, biodiversity indicators mostly relate to pressure on biodiversity, while direct, taxonomic state indicators describing biodiversity remain very rare in the process. The demand for more, structured and direct, taxonomic indicators has already been recognized by e.g. the development of Essential Biodiversity Variables for monitoring biodiversity (Scholes et al., 2012, Pereira et al., 2013) or other policy initiatives like the IUCN red-lists or the conservation status of habitats (Lier et al., 2013). In the case of forests, the taxa at stakes are rarely assessed (e.g. those mostly impacted by forest management, like saproxylic taxa e.g. Paillet et al., 2010). With a lot of pressure and very few biodiversity assessed, it is also likely that the responses in terms of biodiversity preservation remain poorly adapted with the current indicators included in the process. However, classifying not only indicators of criterion 4, but also those implied by other criteria to assess biodiversity, shows that the overall analysis is more valuable that the one based on the dedicated criterion only (Butchart et al., 2010). Although not exhaustive regarding the PSR framework, including indicators from other criteria allows a broader view and a better understanding of the processes at stake regarding biodiversity and opens perspectives of transversal, rather than focused, views.

### Blurred lines between DPSIR categories may cause misinterpretations

As mentioned above, a definition of the indicandum allows to clearly interpret an assessment process (Heink and Kowarik, 2010a). However, we showed that loose definitions of the indicandum blur the visions of the purpose as well as the classifications in the DPSIR framework. Blurred lines exist between several categories: pressure and state, and maybe less evidently, drivers and impacts.

First, in the case of pressure and state indicators assessing sustainable forest management, the process implicitly assumes that if a given indicator belongs to a criterion, then it allows an assessment of at least one component of sustainable forest management: for example, since the indicator dedicated to deadwood (4.5) belongs to the criterion biodiversity, then it is assumed to assess the state of biodiversity. Would this indicator have been included in the criterion 1, it could have been used to assess e.g. the carbon contained in dead forest biomass and soil. However, our approach gives different views that change relatively to the assessment of the state of forests or the biodiversity *per se*. We assumed that biodiversity state indicators should be taxonomic indicators, as using taxonomic data is more precise and specific to assess biodiversity dynamics than proxies (Scholes et al., 2012, Larrieu et al., 2019, Sabatini et al., 2016). Indeed, although most of the time proxies are easier to assess (or sometimes available through other processes like national forest inventories; e.g. Tomppo et al., 2010), they are also less efficient than measuring biodiversity directly. In the case of deadwood, the presence of the habitat (the substrate) does not guarantee the presence of species, since demographic processes may interfere, despite positive correlations between species and habitat (e.g. Lassauce et al., 2011). A colonisation credit may indeed exist (Naaf and Kolk, 2015, Talluto et al., 2017)(ref), and if species sources are absent or too far away to insure effective colonisation, the only presence of a substrate does not indicate properly the state of biodiversity. For these reasons, assuming that indirect (habitat) indicators constitute pressure indicators for biodiversity seems more reasonable. In addition, targeting pressure indicators through dedicated measures (e.g. promoting deadwood in managed forests, or limit its harvesting for woodfuel) rather than action directly involving species populations, notably cryptic taxa such as fungi, bryophytes or lichens, seems more applicable and is more likely to be accepted and understandable by managers or policy makers (Heink and Kowarik, 2010b). This approach notably prevails in the European habitat directive and Natura 2000. Within the PSR framework, it also legitimates the classification and clarifies the policy and management measures and targets. Such measures, in the broader approach of forest assessment, will also improve the overall state of the forest and probably affect more than biodiversity only (e.g. soil fertility).

Second, the case of drivers and impacts is even more blurry and concerns only the assessment of forests within the DPSIR framework. Drivers and impacts represent the two “human” components in the DPSIR framework (response is more related to policy and management measures). As such, drivers and impact are closely linked: for example, we considered roundwood indicator (3.2) as a provisioning indicator, reflecting the wood production service provided by forests, but since it also reflects the wood market and may be modified as such, it may also be considered as a driver indicating the society’s demand for wood products (as several other provisioning indicators, Schwarzbauer and Rametsteiner, 2001). However, since sustainable forest management indicators highlight the current situation rather than the trends, it is more easily understandable as an impact, in terms of ecosystem services. As such, and contrary to the simpler PSR frameworks, the impacts in the DPSIR introduce a notion of value to the system that is useful for communication towards a panel of stakeholders. This aspect is remarkably absent from the simpler PSR framework, but the drivers would very likely be the same (Blauvelt, 2014).

### Limitations of the DSPIR and PSR frameworks and the way forward

The analysis of sustainable forest management (resp. forest biodiversity) indicators through DPSIR (resp. PSR) framework highlights gaps and possible improvements to the sustainable forest management indicator system (Wolfslehner, 2007). However, despite the usefulness of the framework for categorising and analysing the capacity of an indicator system to inform about a subject at stake, the two frameworks we used raise questions and show limitations that are worth mentioning.

The DPSIR framework is essentially anthropocentric and does not clearly allow ecosystem resilience to act as a “response” to a degradation of the state of the system. Indeed, within this framework, a modification of the state of the system has an impact on human well being that is supposed to provoke responses of the society, as shown by the absence of link between pressures and responses in the classical DPSIR scheme (Figure 1). Resilience loop and natural ecosystem dynamics are thus absent from the system, although it includes “doing nothing” as a possible response but still requires a political decision to support it: an example of response could be to let the forest for free development by setting aside forest reserves to improve the state of biodiversity. As such, it reinforces a utilitarian view of nature and ecosystems that was already a point of criticism in the ecosystem services approach, later addressed through the closely related concept of nature contribution to people (McCauley, 2006). In our proposal, we assume that using the classification in ecosystem services to qualify the impact category reinforces this view, but potentially allows one to identify action levers within the society. For example, aesthetic impacts of clearcuts have recently been the subject of debate within the general public as well as specific stakeholder groups (Clausen and Schroeder, 2004, Kearney et al., 2010). Such effect may easily be classified as an impact within the DPSIR framework, and adapted responses may derive from it, as well as indicators to evaluate them. In this example, a multiple response also targeted at the general public (e.g. combining education, limitation of the visual impacts of clearcuts, transition to less intensive management methods such as continuous cover) rather than a single technical one aimed at foresters may be more efficient for public acceptance and overall state of the forests. Conversely, the PSR framework is devoid of a clear human component apart from the policy and management response that may be limitative with respect to an environmental problem, but offers a more straightforward approach for defining and analysing an indicator set, such as we did on biodiversity.

Analysing the links between the categories was beyond the scope of this paper, however it is the next essential step for operationalisation of the approach for sustainable forest management, mostly between those that would be state and pressure indicators (see the attempt by Oettel and Lapin, 2021). Given the relative incompleteness of the system we highlighted, it is likely that links will be hard to assess (see Wolfslehner, 2007 however, but without an implicit classification of the indicators within each category). Although essential, establishing the links between indicators may be purely discursive and theoretical, but may also adopt a correlative (statistical) approach that would be possible with data issued of quinquennal reporting on the State of Europe’s Forests. But, e.g. for biodiversity, it will also require further fundamental research on the functional link between pressures (e.g. substrate availability) and state (diversity of biotic communities), or more generally between indicators and indicandum to validate indicators in a robust way (e.g. Gao et al., 2015, Paillet et al., 2018). For this reason, the approach using an explicit framework may allow to integrate human components and perceptions for a wider identification and validation of indicators (Niemeijer and de Groot, 2008).

## Conclusions

For thirty years now, the quinquenal report on the State of Europe’s Forests has been assessing sustainability in forest management. Thanks to a large panel of indicators, it has been possible to monitor the evolution of different aspects of forest management and policy including production, biodiversity and societal aspects. However, we assumed that this system lacked an analytical framework that would offer a transversal view of forest management and biodiversity issues, as well as help to gain power in terms of assessment and decision making (see Feest, 2013 for a similar essay on EEA indicators), notably since the new definition of sustainable forest management proposed by the European Commission represents a novel and strong boost towards more direct and effective indicators of the state of forest biodiversity (EU Regulation 2020/852). By classifying all the indicators in the DPSIR framework, and biodiversity indicators in the PSR framework, we highlighted the crucial step in the definition of the indicandum, but also the need to use explicit classifications and questions (Svarstad et al., 2008). As trivial as it seems, these steps are often forgotten or implicit in indicator definition and validation processes (Heink and Kowarik, 2010b). In addition, we showed that there are gaps in the indicators sets, notably concerning taxonomic indicators to assess biodiversity dynamics directly, and not only through proxies. By completing these gaps and analysing the links between the different indicators categories, the overall evaluation would gain in efficiency and provide more adapted responses that would allow improving sustainable forest management at the large scale. This would require a strong collaboration between stakeholders, policy makers, and scientists to define and validate indicators. Accordingly, aligning the DSPIR classification of sustainable forest management indicators with available information from forest statistical sources such as National Forest Inventories, will allow creating a baseline for assessing historical and current trends as well as the corresponding effectiveness of the adopted forest management actions at large (e.g. EU-Countries) geographical scales (Bowditch et al., 2020). Such analysis is a step towards more integration and should be used to prioritize actions in favor of sustainability. However, neither sustainable forest management indicators, nor any analytic framework will help solving conflicts without more knowledge on the link between the different categories of the classification to support decision-making processes.

## CRediT authorship contribution statement

Y. Paillet, J. Dorioz, M. Gosselin, J. Marsaud: Conceptualization, Formal analysis; Y. Paillet: Writing - original draft; All the authors: Writing - review & editing.

## Acknowledgements

This paper is issued of different workshops held in Montargis, France (2011) and through the framework of the CHIFFRE project (“Construction Historique des Indicateurs Faune et Flore et Représentations de l’Environnement”, coordinator: G. Bouleau, Irstea, 2012-2014). It also benefited from discussions within the framework of the COST action CA18207 “Biodiversity Of Temperate forest Taxa Orienting Management Sustainability by Unifying Perspectives” (https://www.bottoms-up.eu/). We thank Lisa Paix for her significant kick-off for the finalization of this paper.

